# Cellular transformation by combined lineage conversion and oncogene expression

**DOI:** 10.1101/525600

**Authors:** Biswajyoti Sahu, Päivi Pihlajamaa, Kaiyang Zhang, Kimmo Palin, Saija Ahonen, Alejandra Cervera, Ari Ristimäki, Lauri A. Aaltonen, Sampsa Hautaniemi, Jussi Taipale

**Author notes:** Corresponding author: Jussi Taipale.

## Abstract

Cancer is the most complex genetic disease known, with mutations implicated in more than 250 genes. However, it is still elusive which specific mutations found in human patients lead to tumorigenesis. Here we show that a combination of oncogenes that is characteristic of liver cancer (CTNNB1, TERT, MYC) induces senescence in human fibroblasts and primary hepatocytes. However, reprogramming fibroblasts to a liver progenitor fate, induced hepatocytes (iHeps), makes them sensitive to transformation by the same oncogenes. The transformed iHeps are highly proliferative, tumorigenic in nude mice, and bear gene expression signatures of liver cancer. These results show that tumorigenesis is triggered by a combination of three elements: the set of driver mutations, the cellular lineage, and the state of differentiation of the cells along the lineage. Our results provide direct support for the role of cell identity as a key determinant in transformation, and establish a paradigm for studying the dynamic role of oncogenic drivers in human tumorigenesis.

## Introduction

Cancer genetics and genomics have identified a large number of genes implicated in human cancer (Alexandrov *et al*, 2013, Forbes *et al*, 2017, Garraway *et al*, 2013, Vogelstein *et al*, 2013). Although some genes such as *p53* and *PTEN* are commonly mutated in many different types of cancer, most cancer genes are more lineage-specific. It is well established that human cells are harder to transform than rodent cells (Boehm *et al*, 2005, Chaffer *et al*, 2015, Kamijo *et al*, 1997, Metz *et al*, 1995, Rangarajan *et al*, 2004, Ruley, 1983, Stevenson *et al*, 1986), which can be transformed using only MYC and RAS oncogenes (Land *et al*, 1983, Shih *et al*, 1981, Sinn *et al*, 1987). Seminal experiments by Hahn and Weinberg established already 20 years ago that different human cell types can be transformed using a set of oncogenes that includes the powerful viral large-T and small-T oncoproteins from the SV40 virus (Hahn *et al*, 1999). Despite this early major advance, determining which specific mutations found in human patients lead to tumorigenesis has proven to be exceptionally difficult. This is because although viral oncoproteins are linked to several cancer types (Moore *et al*, 2010), most major forms of human cancer result from mutations affecting tumor-type specific sets of endogenous proto-oncogenes and tumor-suppressors (Haigis *et al*, 2019).

The idea that distinct cellular states promote tumorigenesis is well established in animal models. Many tumor promoting agents (Yamagiwa & Ichikawa, 1918) are not efficient mutagens (reviewed in Diamond *et al*, 1980), suggesting an indirect or epigenetic mechanism for their action. For example, wounding promotes tumorigenesis (Dvorak, 1986), and oncogene activation in combination with a wound environment initiates epidermal tumorigenesis from mouse keratinocytes (Kasper *et al*, 2011). Furthermore, experiments in cultured cells have established that not all oncogenes can transform rodent fibroblasts (Barr, 1998, Daley *et al*, 1987), indicating that at least a subset of oncogenes are lineage-specific. Furthermore, previous studies using genetically modified mouse models have suggested that the oncogenes promote tumorigenesis in a tissue- and context-specific manner. For example, Myc expression in mouse hepatocytes during embryonic development resulted in immediate onset of tumor growth, whereas adult mice developed tumors only after prolonged latency (Beer *et al*, 2004). Similarly, mutant KRAS-G12V is sufficient to induce pancreatic ductal adenocarcinoma in mice, when expressed in embryonic cells of acinar lineage, whereas chronic inflammation in combination with KRAS-G12V expression is required for pancreatic tumorigenesis in adult mice (Guerra *et al*, 2007). In lung, however, the expression of mutant KRAS alone is sufficient for tumor development also in adult mice (Jackson *et al*, 2001; Johnson *et al*, 2001). Taken together, data from experimental animal studies suggests that oncogenes are lineage-specific. However, rodent cells are much easier to transform than human cells, and it is presently not clear whether this is because human cells require more mutagenic hits, or whether a smaller fraction of human cells are susceptible to transformation, resulting in differences in interactions between cellular lineage and transformation between humans and mice. Consistent with the latter possibility, although mutation of the same oncogene or tumor suppressor often causes tumors in similar tissues in mice and humans, also major differences exist. For example, germline loss of one allele of APC leads primarily to small intestinal polyps in mice, but colon cancer in humans; should small intestinal polyps be as easily formed in humans than in mice, small intestinal cancer would be one of the most common cancer types in humans, suggesting that differences in lineage restriction of tumorigenesis play a role in differences of tumor incidence between species.

Despite decades of work, it is still elusive why oncogenes are lineage-specific in humans, and what makes human cells more resistant to oncogenic transformation compared to rodent cells. One possibility is that cell lineage-specific factors could somehow interact with oncogenes to drive most cases of human cancer, and that this process could be at least in some cases specific to human cells, confounding mechanistic studies utilizing simple model cell types and cells from model organisms. Thus, in addition to studying tumorigenesis *in vivo* in model organisms, complementary studies of tumorigenic processes in human cells are critical for fully understanding cancer. However, experimental studies of transforming human cells in their natural environment are clearly neither possible nor ethical. In principle, individual driver genes and their combinations could be identified and validated using particular primary cell types. Such an approach is limited by the fact that for most tissues, sufficient amounts of live human tissue material are hard to obtain. Furthermore, the cell type of origin for most cancer types is not known, and it is commonly assumed that tumors originate from rare and hard-to-isolate subpopulations of cells (e.g. stem cells, or transient progenitor cells in the case of pediatric tumors). Furthermore, although some previous studies have reported oncogene combinations that can transform primary human cells (Drost *et al*, 2015, Matano *et al*, 2015, Park *et al*, 2018), many primary cells may not be at the specific differentiated state that is required for transformation.

These considerations prompted us to systematically investigate the factors required for transformation of human cells using a combination of cell fate conversion and oncogene activation. This approach has the potential to overcome the limitations inherent to experimental animal models and primary human cells. Importantly, our approach recapitulates the difficulty of transforming human cells, creating a platform for detailed studies for the interplay of cell identity and epigenetic state with oncogenic drivers in human cells.

## Results

### Generating proliferative induced hepatocytes using defined transcription factors and oncogenic drivers

Many human cell types can be converted to other cell types via a pluripotent state (Takahashi *et al*, 2007). However, as pluripotent cells are tumorigenic in nude mice, we chose to use direct lineage conversion (Davis *et al*, 1987, Pang *et al*, 2011, Sekiya *et al*, 2011) in combination with oncogene expression to identify the set of factors that define a particular type of human cancer cell. For this purpose, we developed a cellular transformation assay protocol, in which human fibroblasts (HF) are converted to induced hepatocytes (iHeps) using lentiviral expression of a combination of lineage-specific transcription factors (TF), followed by ectopic expression of liver cancer-specific oncogenes (Fig. 1A). Transdifferentiation of fibroblasts to iHeps has previously been reported by several groups (Du *et al*, 2014, Huang *et al*, 2014, Morris *et al*, 2014, Sekiya *et al*, 2011). To identify an optimal protocol for generating iHeps from HFs (from human foreskin), we tested the previously reported combinations of TFs in parallel transdifferentiation experiments and analyzed the efficiency of iHep conversion by measuring the mRNA levels for liver markers (Du *et al*, 2014, Huang *et al*, 2014, Morris *et al*, 2014) such as *ALBUMIN, TRANSFERRIN,* and *SERPINA1* at different time points (Fig. 1B, **Fig. S1**). The combination of three TFs, HNF1A, HNF4A and FOXA3 (Huang *et al*, 2014) resulted in the most efficient iHep generation, based on the observation that out of all combinations tested, this combination resulted in the highest expression level of liver-specific genes at two, three, and four weeks after iHep induction (Fig. 1B). This protocol also resulted in most efficient lineage conversion based on the analysis of cell morphology; by two weeks after iHep induction, the cells lost their fibroblast phenotype and formed spherical iHep progenitor colonies, from which immature, proliferative iHeps migrated outward. The iHeps fully matured to non-proliferative iHeps by six to seven weeks after induction (Fig. 2A and **Fig. S2**).

**Figure 1.**
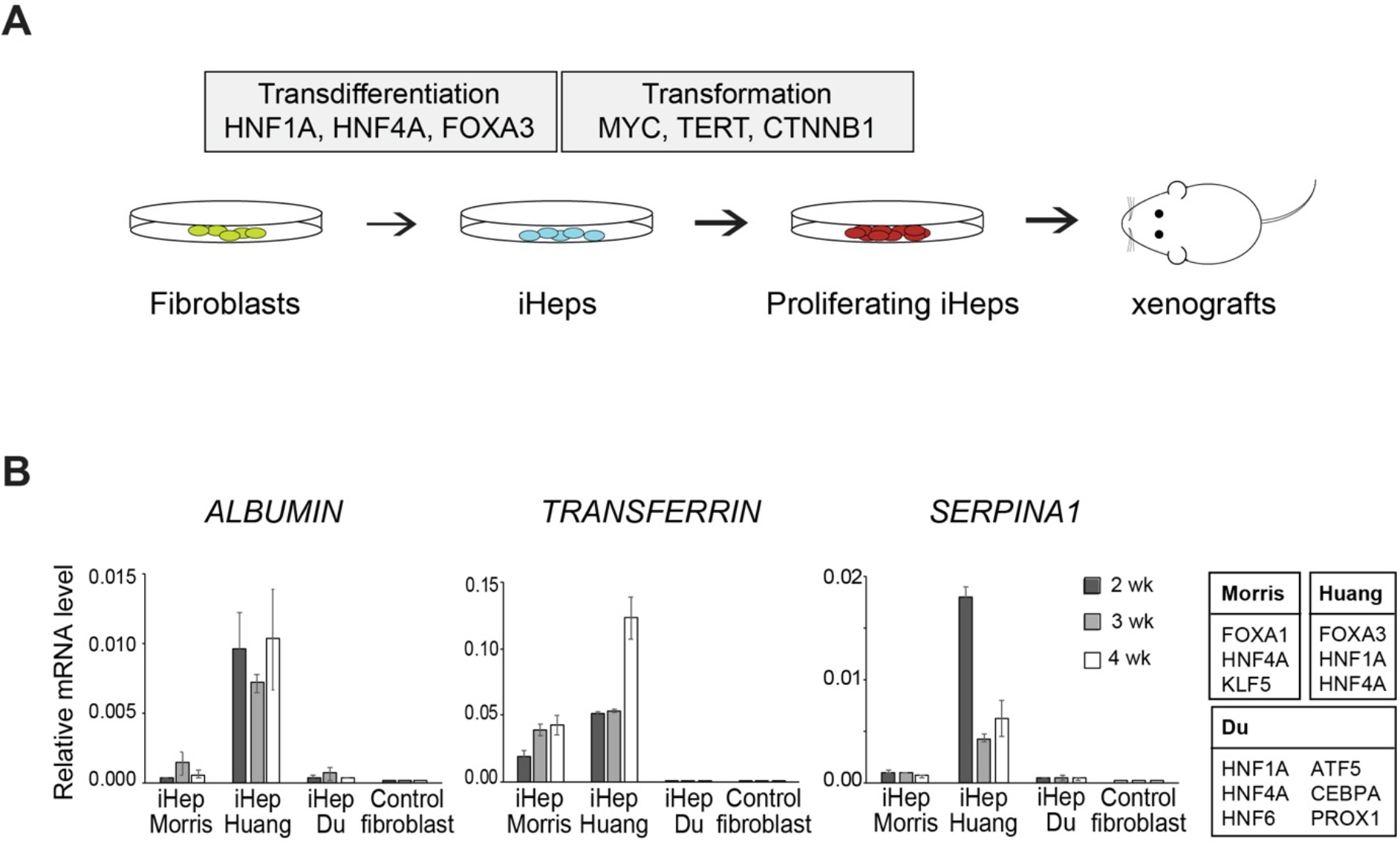
Generating proliferative induced hepatocytes using defined transcription factors and oncogenic drivers. (**A**) Schematic outline of the cell transformation assay for making lineage-specific cancer by lentiviral expression of three lineage-specific TFs to convert HFs to induced hepatocytes (iHep) and defined oncogenic drivers to transform iHeps to proliferating and tumorigenic cells. (**B**) Comparison of TF combinations (Du *et al*, 2014, Huang *et al*, 2014, Morris *et al*, 2014) for converting human fibroblasts to iHeps by detecting transcript levels for liver marker genes (*ALBUMIN*, *TRANSFERRIN* and *SERPINA1/α-1-antitrypsin*) by qRT-PCR at different time points after iHep conversion, normalized to GAPDH levels (mean ± standard error).

**Figure 2.**
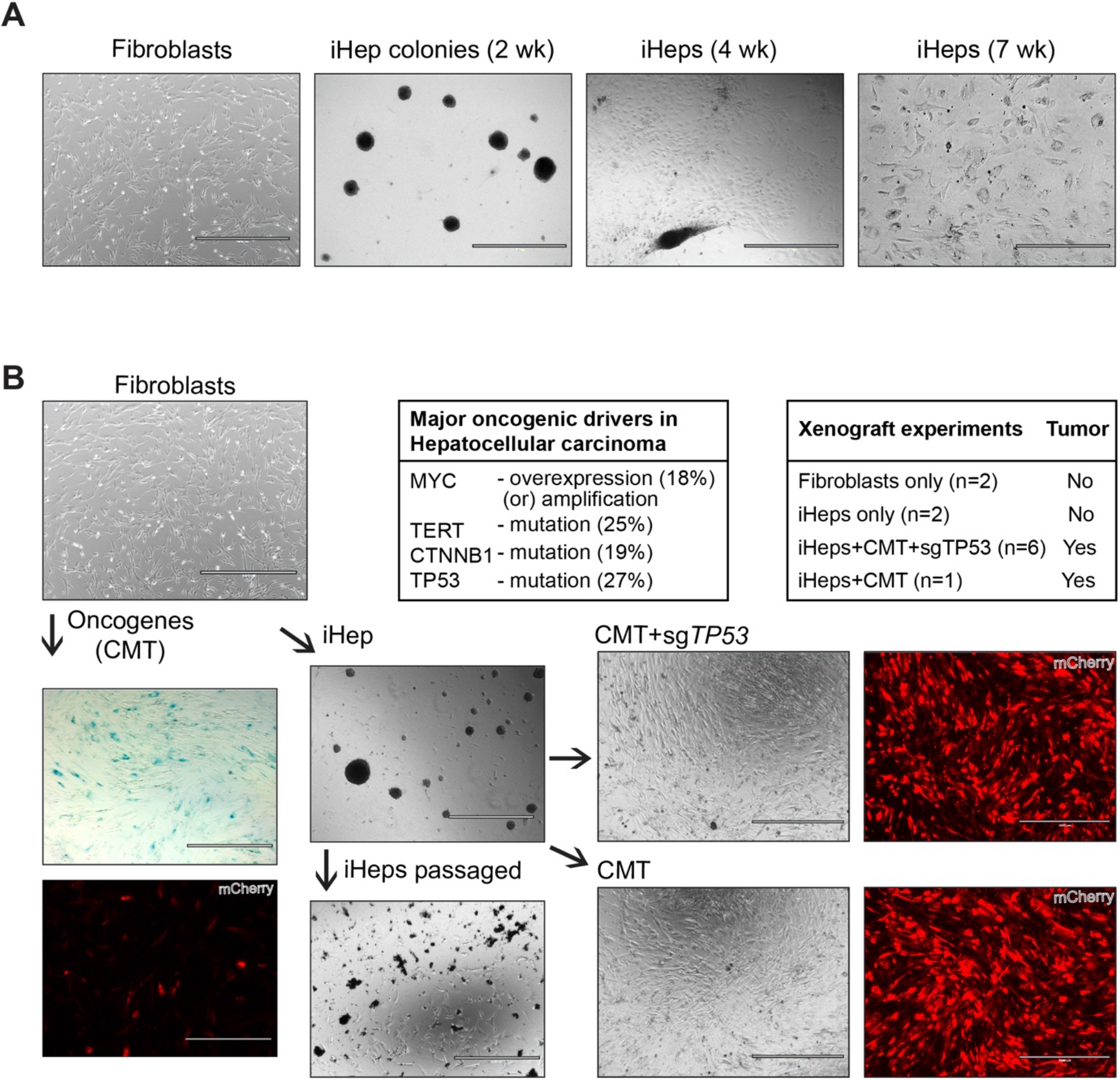
Oncogene exposure transforms induced hepatocytes but not control fibroblasts. (**A**) Phase contrast microscope images showing the phenotype and morphology of the cells in the course of conversion of fibroblasts to iHeps at different times points after transduction with a cocktail of three TFs HNF1A, HNF4A and FOXA3 (Huang *et al*, 2014). (**B**) Generation of highly proliferative iHep cells by transducing iHeps with two pools of liver cancer-specific oncogenic drivers, a list of xenograft experiments in nude mice that were used to test the tumorigenicity of different conditions, and mutation rates of the oncogenic drivers as reported in the COSMIC database for HCC and MYC amplification as reported in (Cancer Genome Atlas Research Network, 2017). CMT pool contains three oncogenes CTNNB1^T41A^, MYC, and TERT, and CMT+sg*TP53* pool contains the same oncogenes along with constructs for *TP53* inactivation by CRISPR-Cas9. Phase contrast microscope images showing the phenotype and morphology of the cells. Oncogenes are co-transduced with fluorescent reporter mCherry for detection of transduced cells. Oncogene transduction to fibroblasts fails to transform the cells, passaging of oncogene-expressing fibroblasts results in cellular senescence as demonstrated by beta-galactosidase staining and loss of mCherry-positive oncogene-expressing cells from the fibroblast population. Passaging of iHeps without oncogenes results in apoptosis after few passages. Scale bar 1000 μm unless otherwise specified.

### Oncogene exposure transforms induced hepatocytes but not control fibroblasts

To determine whether the iHeps could be transformed to liver cancer-like cells, we first plated immature (1 to 3 weeks post-transdifferentiation) iHeps on collagen-coated dishes and maintained them in hepatocyte culture media (HCM). Under such conditions, the iHeps mature, and their proliferation is arrested after two to three passages (Huang *et al*, 2014); after this point, further passaging induces cell death (Fig. 2B). To confer the immature iHeps with unlimited proliferative potential and to drive them towards tumorigenesis, we transduced them with a set of the most common driver genes for liver cancer using lentiviral constructs. For this purpose, we chose the five oncogenic drivers with the highest number of recurrent genetic alterations reported for liver cancer or hepatocellular carcinoma (HCC; from COSMIC, https://cancer.sanger.ac.uk/cosmic); these included four oncogenes, telomerase (*TERT*), β-catenin (*CTNNB1*), PI3 kinase (*PIK3CA*), and the transcription factor NRF2 (*NFE2L2*), as well as one tumor suppressor, p53 (*TP53*). In addition, we included the oncogene *MYC*, which is under tight control in normal cells (Lowe *et al*, 2004), but overexpressed in many cancer types, including HCC (Kalkat *et al*, 2017). Lentiviral expression of the fluorescent reporter mCherry with the oncogenic drivers in different combinations revealed that the pool of three oncogenes, *i.e.* constitutively active β-catenin (CTNNB1^T41A^), MYC and TERT, together with *TP53* inactivation by CRISPR-Cas9 (CMT+sg*TP53*) resulted in highly proliferative iHeps with apparently unlimited proliferative potential (> 50 passages over more than one year; Fig. 2B). Transduction of iHeps with other combinations of the oncogenic drivers did not result in sustained proliferative phenotype, indicating that the iHeps are not susceptible to any oncogenic insult but to a specific combination of these three oncogenes. Importantly, expression of the three oncogenes CTNNB1^T41A^, MYC and TERT (CMT) alone also resulted in similar iHeps with long-term proliferative potential (Fig. 2B). By contrast, ectopic expression of these oncogenic drivers in HFs failed to yield transformed, proliferating fibroblasts, and rather resulted in cellular senescence and loss of the oncogene-transduced cells from the fibroblast population (Fig. 2B). This is the first instance to our knowledge where HFs can be directly transformed using this minimal combination of defined factors, indicating that lineage-specific TFs are the missing link for human cellular transformation using oncogenic drivers.

### Tumorigenic properties of the transformed iHeps

To test for the tumorigenicity of the proliferative iHeps, we performed xenograft experiments. Subcutaneous injection of the CMT+sg*TP53* transformed iHeps, but not the control fibroblasts or iHeps with lineage-specific TFs alone, into nude mice resulted in tumor formation (Fig. 3A and B). The process was reproducible in subsequent experiments; in addition, the effect was not specific to the fibroblast line used, as we also successfully reprogrammed another HF cell line (human fetal lung fibroblast) using the same lineage-specific TFs, and transformed it using the same set of oncogenic drivers. The xenograft tumors from the CMT+sg*TP53* transformed iHeps derived from either fibroblast line can be detected by *in vivo* fluorescent imaging as early as 11-12 weeks (Fig. 3B). Importantly, the histology of CMT+sg*TP53* tumors harvested at 20 weeks show highly malignant and proliferative features (Fig. 3C). Similarly, the CMT-transformed iHeps without *TP53* inactivation also resulted in tumor formation in nude mice 12 weeks post-injection (Fig. 3B). These results demonstrate that both CMT and CMT+sg*TP53* transformed iHeps are tumorigenic, and indicate that ectopic expression of defined lineage-specific TFs and oncogenes can reprogram and transform HFs into cells that can robustly initiate tumors in nude mice.

**Figure 3.**
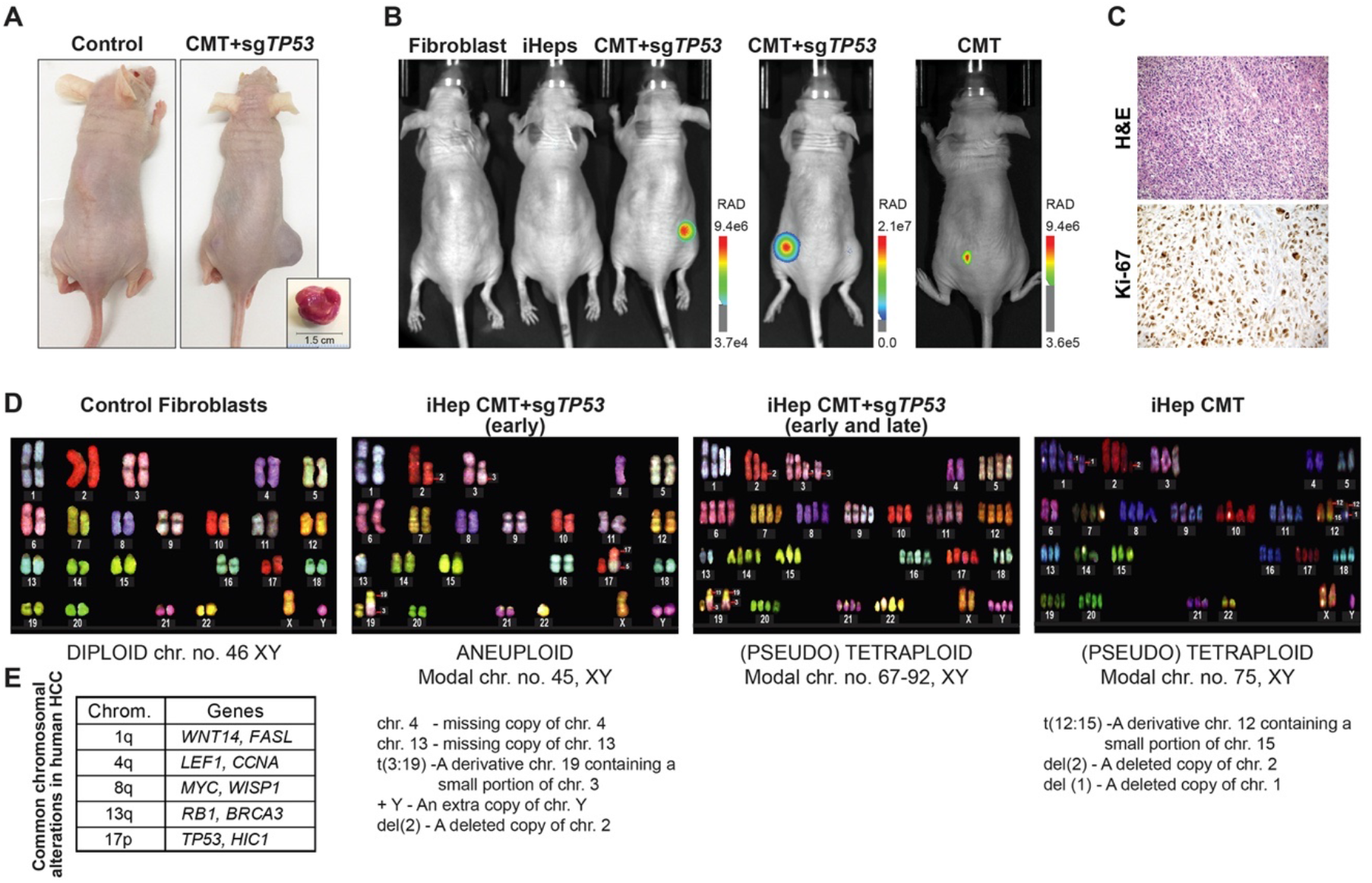
Tumorigenic properties of the transformed iHeps. (**A**) Subcutaneous injection of transformed iHeps results in xenograft tumors in nude mice (tumor size of 1.5 cm ∼ 23 weeks after xenotransplantation). Proliferative iHeps transduced with defined CMT oncogenes with *TP53* inactivation (CMT+sg*TP53*) or control iHeps without oncogenes were used in the injections. (**B**) *In vivo* imaging of xenograft tumors ∼12 weeks after implantation. Two biological replicate experiments are shown for CMT+sg*TP53* cells with iHep conversion and oncogene transduction with *TP53* inactivation performed in two separate human fibroblast cell lines (foreskin fibroblast [*left panel*] and fetal lung fibroblast [*middle*]) as well as proliferative CMT iHeps without *TP53* inactivation (*right*). Fluorescence signal emitted by mCherry co-transduced with the oncogenes is detected *in vivo* using the Lago system (scale bar = radiance units). Control mice are injected with either fibroblasts or iHeps. (**C**) Histological analysis of CMT+sg*TP53* tumor tissue harvested at 20 weeks. Hematoxylin-eosin (H&E) staining for general histology and immunohistochemical staining for Ki-67 for cell proliferation (100x magnification). Note that the appearance of the tumor is consistent with both poorly differentiated hepatic tumor (WHO Classification of tumors, 5^th^ edition, 2019) or sarcoma. Differential diagnosis from sarcoma is accomplished by analysis of marker gene expression (see Fig. 5B). (**D**) Analysis of chromosomal aberrations in the transformed iHeps by spectral karyotyping. CMT+sg*TP53* cells were analyzed at passage 18 (*early*) and passage 50 (*late*) and CMT cells at passage 18. Fibroblasts have normal diploid karyotype (46, XY, representative spectral image on *left*) and transformed iHeps show aneuploidies as indicated in the figure. Early passage CMT+sg*TP53* cells show two different populations with two distinct modal chromosome numbers (45, XY and 67-92, XY, representative spectral image for 45, XY on *middle-left*). Late passage CMT+sg*TP53* cells have modal chromosome number 67-92, XY (*middle-right*) and CMT cells 75, XY (*right*). In the text box below the images, recurrent chromosomal aberrations seen in majority (>90%) of the cells are reported. (**E**) Frequencies of chromosomal alterations reported for human HCC samples [see (Moinzadeh *et al*, 2005)].

Cancer genomes harbor large-scale chromosomal aberrations and are characterized by aneuploidy (Palin *et al*, 2018, Taylor *et al*, 2018). To understand the gross chromosomal aberrations in the transformed tumorigenic CMT and CMT+sg*TP53* iHeps compared to normal HFs, we performed spectral karyotyping, which showed a normal diploid male (46, XY) in HFs and aneuploid karyotypes in transformed iHeps (Fig. 3D). The aneuploid transformed iHeps with CMT+sg*TP53* at early passage were characterized by two different populations with two distinct modal chromosome numbers (Fig. 3D). The modal chromosome number of the first population was 45, XY, whereas the second population was pseudotetraploid, with a modal chromosome number between 67-92, XY; this pseudotetraploid state was consistently observed in late passage transformed iHeps. The major chromosomal aberrations that were similar between the two populations were missing copies of chromosomes 4 and 13, a derivative of chromosome 19 containing a small portion of chromosome 3 [t3:19], an extra copy of Y and a loss of most of the p arm of chromosome In comparison, the most common chromosomal aberrations reported in HCC are the gains of 1q (suggested target genes include *WNT14, FASL*) and 8q (*MYC, WISP1*) and the loss of 17p (*TP53, HIC1*), followed by losses of 4q (*LEF1, CCNA*) and 13q (*RB1, BRCA3*) (Moinzadeh *et al*, 2005, Cancer Genome Atlas Research Network, 2017) (Fig. 3E). The first three chromosomal aberrations are expected not to be present in our case, as the transformation protocol leads to activation of the Wnt pathway and MYC expression, and loss of p53. Consistently with this, we did not observe lesions in 1q, 8q or 17p in our cells. However, other common aberrations found in HCC cells, loss of chromosomes 4 and 13 were detected in our transformed CMT+sg*TP53* iHep cells (Fig. 3D and E). However, these chromosomal aberrations appeared not to be necessary for formation of tumors, as in the absence of targeted loss of p53 in CMT iHep cells, we did not observe these lesions (Fig. 3D). However, both CMT+sg*TP53* and CMT iHeps displayed pseudotetraploidy, similar to what is observed in about 25% of the HCC cases, especially in the highly proliferative cases with poor prognosis (Bou-Nader *et al*, 2020) (Fig. 3D and E). These results indicate that the transformed iHeps have similar chromosomal aberrations to those reported earlier in liver cancer, consistent with their identity as HCC-like cells.

### Dynamic activity of oncogenes during tumorigenesis

To understand the gene expression dynamics and to map the early events of lineage conversion and oncogenic transformation, we performed single cell RNA-sequencing (scRNA-seq) of HFs and iHeps with or without oncogene transduction. The cells were clustered according to their expression profiles using Seurat (Satija *et al*, 2015); a total of 15 separate clusters of cells were identified during the course of the transdifferentiation and reprogramming and visualized by t-distributed stochastic neighbour embedding (t-SNE) plots (van der Maaten *et al*, 2008) (Fig. 4A and B). Importantly, the scRNA-seq indicated that the CMT-transformed iHeps are a clearly distinct population of cells compared to the iHeps, whereas CMT-transduced HFs are more similar to the control HFs (Fig. 4B).

**Figure 4.**
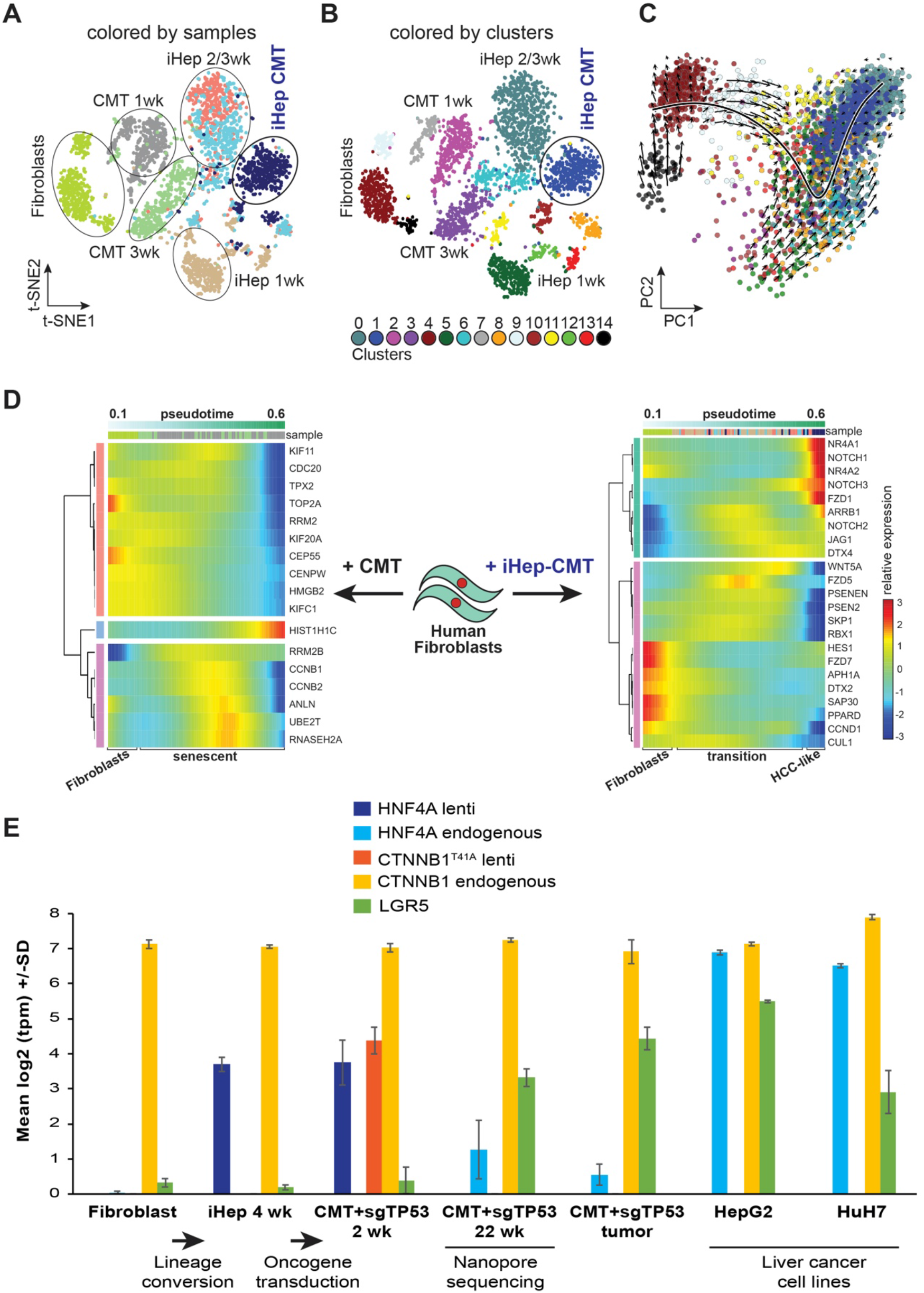
Dynamic activity of oncogenes during tumorigenesis. (**A, B**) t-SNE plots of 3,500 single cells from fibroblasts, iHeps at one–three weeks after iHep induction, iHeps transduced with CMT oncogenes at one week and harvested for scRNA-seq two weeks later, and fibroblasts transduced with CMT oncogenes and harvested at one and three weeks. Cells are colored by sample (**A**), and distinct clusters (**B**) based on their expression profiles with sample collection time points indicated. (**C**) Principal component analysis (PCA) projection of single cells from control fibroblasts, iHeps at one–three weeks after iHep induction, and CMT-iHeps two weeks after oncogenes shown with velocity field with the observed states of the cells shown as circles and the extrapolated future states shown with arrows for the first two principal components. Cells are colored by cluster identities corresponding to Fig. 4B. (**D**) Relative expression of the genes from the Notch signaling pathway (*panel on the right*) across pseudotime in the single-cell RNA-seq data from control fibroblasts, iHeps at one– three weeks after iHep induction, and CMT-iHeps two weeks after oncogenes (the expression of a gene in a particular cell relative to the average expression of that gene across all cells). Relative expression of the senescence marker genes (Marthandan *et al*, 2016) (*panel on the left*) from control fibroblasts and fibroblasts transduced with CMT oncogenes and harvested at one and three weeks after transduction. Color codes illustrating sample and cluster identities correspond to the colors in Fig. 4A and B, respectively. (**E**) Expression levels [log2(transcripts per million, tpm)] for *LGR5* as well as lentiviral and endogenous *HNF4A* and *CTNNB1* in bulk RNA-seq measurements from control fibroblasts, iHeps at four weeks of differentiation, CMT+sg*TP53*-iHeps at two and 22 weeks after oncogene transduction, xenograft tumor from CMT+sg*TP53* cells, and from liver cancer cell lines HepG2 and HuH7 (mean ±standard error, n=3). Nanopore sequencing was performed from the CMT+sg*TP53* cells at 22 weeks after oncogene transduction as indicated in the figure and used for identifying the genomic insertions of the lentiviral constructs (**Table S1**).

To determine the trajectory of differentiation of the cells, we performed RNA velocity analysis (La Manno *et al*, 2018), which determines the direction of differentiation of individual cells based on comparison of levels of spliced mRNAs (current state) with nascent unspliced mRNAs (representative of future state). This analysis confirmed that the cell populations analyzed were differentiating along the fibroblasts–iHep–transformed iHep axis (Fig. 4C). We next identified marker genes for each cell cluster (see **Materials and methods**). This analysis revealed that CMT-iHeps have a distinct gene expression signature and that they have lost the fibroblast gene expression program during the course of the reprogramming (**Fig. S3**). These results indicate that the iHep conversion and transformation have led to generation of liver-cell like transformed cells.

To further analyze gene expression changes during reprogramming and transformation, we performed pseudo-temporal ordering analysis of the scRNA-seq (see **Materials and methods**). Consistently with the RNA velocity analysis, the pseudotime analysis showed transition from fibroblasts to iHeps and subsequently to CMT-transformed iHeps (**Fig. S4**). Similarly, CMT-transduced HF cells were ordered across pseudo-temporal timeline (**Fig. S4**). The scRNA-seq analyses allow detection of the precise early events that occur during iHep formation and the origin of HCC by mapping the gene expression changes in the cells across the pseudotime. Furthermore, analyzing the molecular changes upon CMT-transduction provided mechanistic understanding of why oncogenes fail to transform HFs without iHep conversion (Fig. 4D); the pseudotime analysis of gene expression changes from CMT-transduced HFs at one and three weeks compared to control HFs was highly similar to the previously reported signature of cellular senescence [17 out of 18 genes (Marthandan *et al*, 2016)] (Fig. 4D). The senescence signature was much weaker both during transdifferentiation of the iHeps and during their transformation (**Fig. S5)**; instead, during iHep differentiation, the expression of non-canonical Wnt pathway components, including Wnt5a ligand and the Frizzled 5 receptor, were upregulated (Fig. 4D). During transformation, the exogenous CTNNB1^T41A^ activated the canonical Wnt pathway, suppressing expression of the non-canonical ligand Wnt5a. We also observe activation of the NOTCH pathway early during tumorigenesis; expression of *NOTCH1*, *NOTCH3* and their ligand *JAG1* (Fig. 4D, top) are strongly upregulated, together with the canonical NOTCH target gene *HES* (Borggrefe *et al*, 2009) and the liver specific target *NR4A2* (Zhu *et al*, 2017). These results are consistent with the proposed role of the NOTCH pathway in liver tumorigenesis (Villanueva *et al*, 2012, Zhu *et al*, 2017).

To map the temporal dynamics of expression of the introduced transgenes and endogenous genes, we performed bulk RNA-seq analysis from the iHeps and tumorigenic CMT+sg*TP53* iHeps that were used for the xenograft implantation, as well as cells derived from the resulting tumors. We also mapped the copy-numbers of the lentiviral transgenes in the CMT+sg*TP53* iHeps using Nanopore sequencing (see **Methods**). This analysis revealed that the iHeps initially expressed all the transgenes, but that during transformation, a clonal cell line that had lost HNF4A and CTNNB1 insertions was selected (**Table S1**). Consistent with reprogramming to liver and tumor cells, respectively, the clonal line expressed endogenous HNF4A, and high levels of the Wnt pathway modulator and stem cell marker LGR5 (Fig. 4E). In summary, we find here that CMT transduced HFs undergo senescence, whereas the proliferative phenotype of CMT-iHep cells is associated with dynamic gene expression changes that affect the Wnt and NOTCH signaling pathways.

### Direct conversion of human fibroblasts to liver cancer cells results in up-regulation of liver cancer markers

To determine the changes in gene expression and chromatin accessibility in the proliferative iHeps, we first performed bulk RNA-seq analysis from the tumorigenic CMT and CMT+sg*TP53* iHeps that were used for the xenograft implantation, as well as cells derived from the resulting tumors. Importantly, the genes that were differentially expressed in both CMT-and CMT+sg*TP53*-transformed iHeps compared to fibroblasts showed a clear and significant positive enrichment for the previously reported “subclass 2” liver cancer signature (Hoshida *et al*, 2009), associated with proliferation and activation of the MYC and AKT signaling pathways (Fig. 5A). The effect was specific to liver cancer, as we did not observe significant enrichment of gene expression signatures of other cancer types (**Fig. S6**). During the reprogramming, we observed a clear up-regulation of common liver marker genes such as *ALB*, *APOA2*, *SERPINA1*, and *TF*, and down-regulation of fibroblast markers such as *MMP3*, *FGF7*, *THY1*, and *FAP*, in proliferative and tumorigenic iHeps. Importantly, the xenograft tumor from the CMT+sg*TP53* cells retained similar liver-specific gene expression profile (Fig. 5B). We also detected a clear up-regulation of several liver cancer marker genes such as *AFP, GPC3, SAA1,* and *VIL1* in transformed iHeps and in CMT+sg*TP53* tumors compared to control fibroblasts (Fig. 5B); *AFP* was also found among the most enriched genes (**Fig. S7**) in both CMT+sg*TP53-* and CMT-transformed iHeps. Furthermore, we observed a negative correlation between the CMT+sg*TP53* and CMT iHep specific genes and the genes positively associated with liver cancer survival (**Fig. S8**), lending further credence to liver cancer-identity of the CMT+sg*TP53* and CMT transformed iHeps.

**Figure 5.**
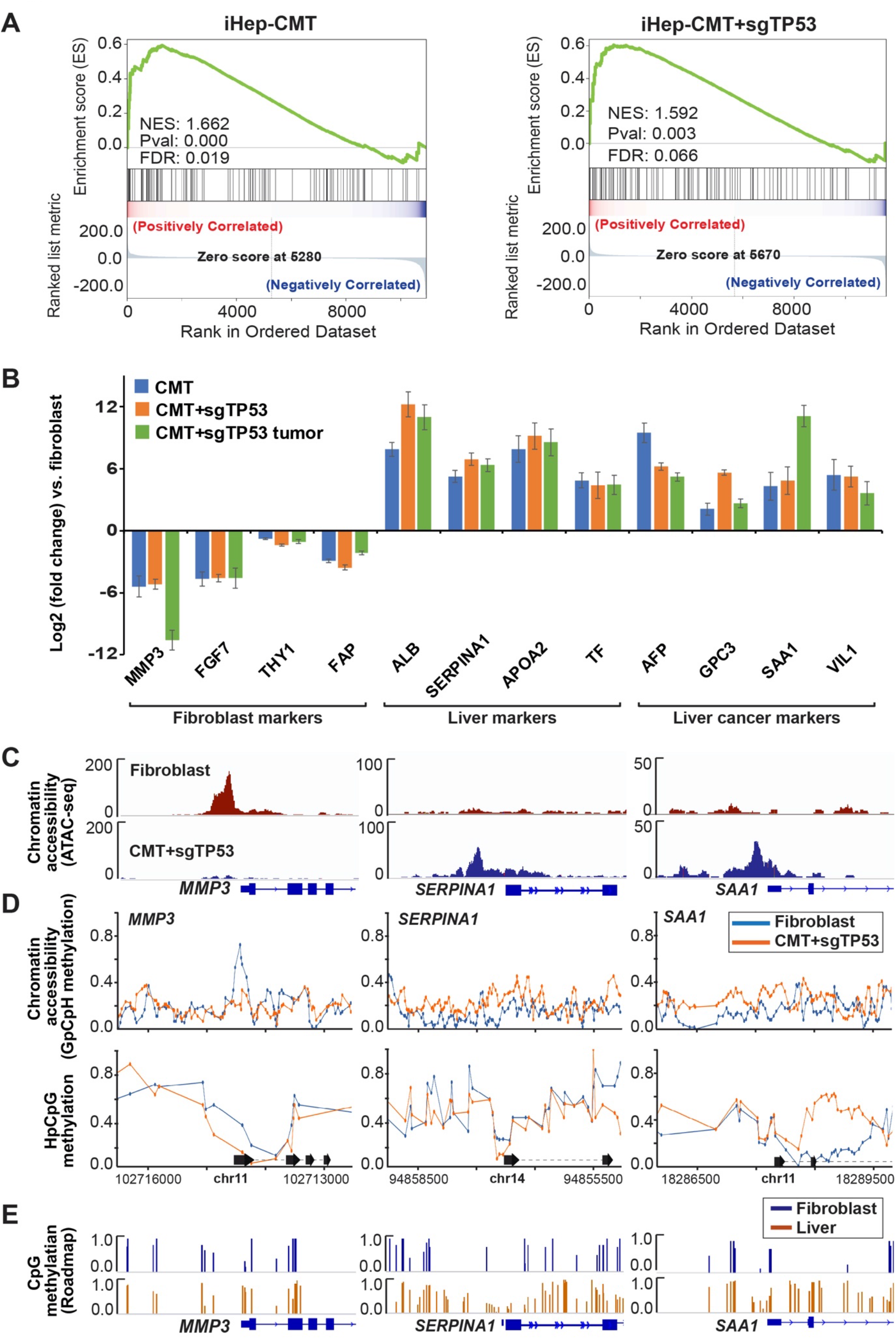
Transformed iHeps show gene expression profile similar to liver cancer. (**A**) Gene set enrichment analysis (GSEA) results for CMT-iHeps and CMT+sg*TP53*-iHeps compared to control fibroblasts against liver cancer signature [Subclass 2 (Hoshida *et al*, 2009)] from molecular signatures database (MSigDB). Positive normalized enrichment score (NES) reflects overrepresentation of liver cancer signature genes among the top ranked differentially expressed genes in CMT-iHep and CMT+sg*TP53*-iHep conditions compared to control fibroblasts. (**B**) Differential expression levels [log2(fold change)] of marker genes for fibroblasts, hepatocytes, and liver cancer in bulk RNA-seq measurements from CMT+sg*TP53*-iHeps and CMT-iHeps at p20 (∼22 weeks after oncogene transduction) as well as xenograft tumor from CMT+sg*TP53* cells against control fibroblasts (mean ±standard error, n=3). (**C**) IGV snapshots for promoter regions of representative genes from fibroblast markers (*MMP3*), liver markers (*SERPINA1/α-1-antitrypsin*), and liver cancer markers (*SAA1*) showing ATAC-seq enrichment from fibroblast and CMT+sg*TP53*-iHeps. (**D**) Chromatin accessibility and CpG methylation of DNA measured using NaNoMe-seq. Cytosine methylation detected using Nanopore sequencing from CMT+sg*TP53*-iHeps and control fibroblasts is shown for promoter regions of representative genes from fibroblast markers (*MMP3*), liver markers (*SERPINA1/α-1-antitrypsin*), and liver cancer markers (*SAA1*) using a window of TSS ±1500 bp. GpCpH methylation (all GC sequences where the C is not part of a CG sequence also, top) reports on chromatin accessibility, whereas HpCpG methylation reports on endogenous methylation of cytosines in the CpG context. (**E**) CpG methylation detected using bisulfite-sequencing from primary human foreskin fibroblasts and from normal adult liver [data from the Roadmap Epigenomics Consortium (2015)]. IGV snapshots from the genomic loci corresponding to the *MMP3*, *SERPINA1*, and *SAA1* promoters (same regions as indicated in Fig. 5D) showing methylation proportions [methylated calls / (methylated calls + unmethylated calls)] for all CpGs covered by at least 4 reads.

ATAC-seq analysis of the fibroblasts and CMT+sg*TP53* cells revealed that the changes in marker gene expression were accompanied with robust changes in chromatin accessibility at the corresponding loci (Fig. 5C). To assess chromatin accessibility and DNA methylation at a single-allele level, we performed NaNoMe-seq (see **Materials and methods**), where accessible chromatin is methylated at GpC dinucleotides using the bacterial methylase M.CviPI (Kelly *et al*, 2012). Sequencing of the genome of the treated cells using single-molecule Nanopore sequencer then allows both detection of chromatin accessibility (based on the presence of methylated cytosines at GC dinucleotides) and DNA methylation at CG dinucleotides. This analysis confirmed the changes in DNA accessibility detected using ATAC-seq (Fig. 5D). Changes in DNA methylation at promoters of the differentially expressed genes were relatively minor (Fig. 5D), suggesting that the mechanism of reprogramming does not critically depend on changes in CpG methylation at the marker loci. This is supported by data from primary human cells obtained using bisulfite-sequencing by the Roadmap Epigenomics Consortium (2015); in this dataset as well, only minor differences are observed in the CpG methylation pattern at the marker gene loci between fibroblasts and normal adult liver (Fig. 5E), suggesting that the differences in marker expression are not caused by CpG methylation induced gene silencing. Taken together, these results indicate that our novel cell transformation assay using lineage-specific TFs and cancer-specific oncogenes can reprogram fibroblasts to lineage-specific cancer that bears a gene expression signature similar to that observed in HCC.

### Cellular lineage and the differentiated state of cells along the lineage are critical for tumorigenesis

To identify the necessary and sufficient factors that define lineage-specific cancer types we have here developed a novel cellular transformation protocol, and, for the first time, report direct conversion of HFs to liver cancer cells. First, lentiviral expression of three lineage-specific TFs reprograms HFs to iHeps, and subsequent ectopic expression of liver cancer-specific oncogenic factors transforms iHeps to a highly proliferative and tumorigenic phenotype with chromosomal aberrations and gene expression signature patterns similar to HCC. Based on RNA-seq analysis, ectopic expression of FOXA3, HNF1A, and HNF4A resulted in expression levels in iHeps that are comparable to those observed in liver cancer cell lines (**Fig. S9**). During cellular transdifferentiation and transformation, the expression of HNF1A remains at a relatively constant level, whereas the expression of HNF4A and FOXA3 is lower in the transformed iHeps than in the parental iHeps (**Fig. S9**). Genomic sequences obtained from the NaNoMe-seq experiment revealed that the lentiviral constructs for HNF4A and FOXA3 are no longer present in the iHeps that have reached the highly proliferative and tumorigenic stage (**Table S1**). However, the expression of respective endogenous genes is induced to a level that is enough to sustain the expression of HNF4A target genes (such as *FABP1*, *APOA1,* and *APOB*) as well as known liver marker genes (*AFP*, *ALB*, and *TF*), validating the hepatic identity of the transformed cells (**Fig. S9**).

Importantly, lineage-conversion by specific TFs is required for the transformation process since the same oncogenic drivers alone do not transform HFs (Fig. 6A). After lineage conversion by the defined TFs, oncogenes alone (MYC, CTNNB1 and TERT) are sufficient to drive the transformation with or without inactivation of the tumor suppressor *TP53*. In contrast, oncogene transduction induces senescence in both HFs and in differentiated adult human hepatocytes (Fig. 6A). Interestingly, ectopic expression of TFs with the same vectors as used for iHep conversion (HNF1A, HNF4A, and FOXA3) one week prior to oncogene transduction protects the hepatocytes from oncogene-induced senescence (Fig. 6A). These results show that fully differentiated non-proliferative hepatocytes are not susceptible for transformation by the liver-specific set of oncogenes, suggesting that in addition to cellular lineage, also the differentiated state of cells along the lineage is critical for tumorigenesis. This finding is consistent with our experiments studying transformation in reprogrammed induced neurons (iNs). As these cells become terminally differentiated and post-mitotic immediately upon transdifferentiation, neither medulloblastoma nor neuroblastoma-specific oncogenes were able to make them re-enter the cell cycle (Fig. 6A, **Fig. S10**). These results establish a paradigm for testing the tumorigenicity of combinations of cancer genes, and their interactions with cellular lineage and differentiated state (Fig. 6B). In addition, reprogramming normal cells to cancer cells allow “live” analysis of the early stages of the tumorigenic program, facilitating approaches towards early molecular detection and prevention of cancer.

**Figure 6.**
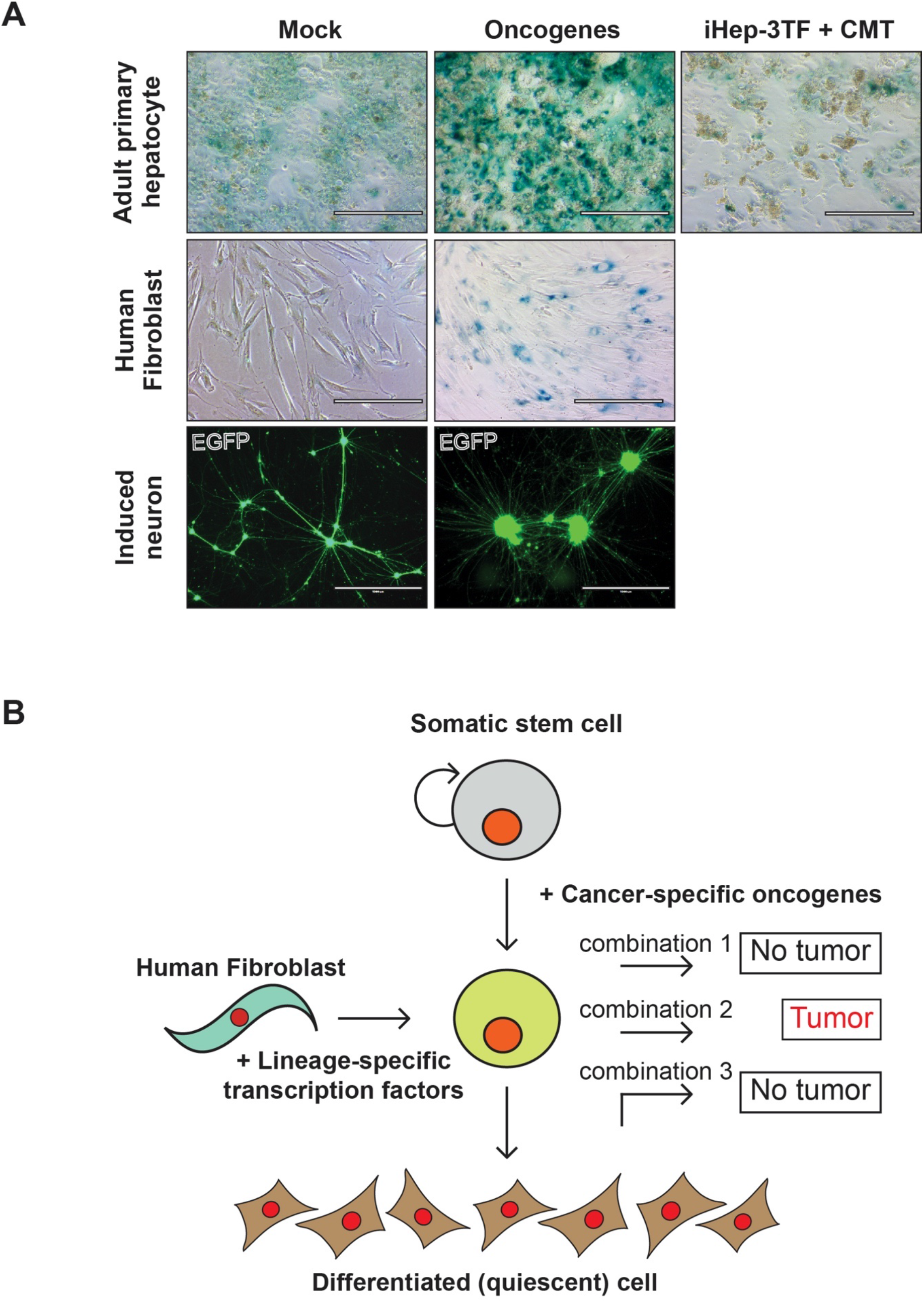
Direct conversion of human fibroblasts to liver cancer cells. (**A**) (*Top*) Beta-galactosidase staining as a marker of cellular senescence in primary human hepatocytes (control), after transduction of CMT oncogenes, or after transduction with iHep-TFs (HNF1A, HNF4A, FOXA3) followed by CMT oncogene transduction one week later (stained three weeks after first transduction). (*Middle*) Beta-galactosidase staining as a marker of cellular senescence in control fibroblasts and fibroblasts transduced with CMT oncogenes and stained at p4. (*Bottom*) Fluorescent microscope images of induced neurons with and without oncogene transduction (at three weeks of neuronal differentiation) visualized using EGFP at ten weeks after neuronal conversion. (**B**) Schematic presentation of the molecular approach for identifying minimal determinants of tumorigenesis in specific tissues. Lineage-specific transcription factors are used to reprogram human fibroblasts to precise cellular identity (*left*), whose transformation by specific combinations of oncogenes (*right*) can then be tested. This approach combined with single-cell RNA-seq and RNA velocity analyses allows also analysis of which cell type along the stem cell to terminally differentiated cell axis (*top to bottom*) is susceptible for transformation.

## Discussion

In the past half-century, a very large number of genetic and genomic studies have been conducted using increasingly powerful technologies, resulting in identification of more than 250 genes that are recurrently mutated in cancer. However, in most cases, the evidence that the mutations in the genes actually cause cancer is correlative in nature, and requires assumptions about background mutation frequency and rates of clonal selection in normal tissues (Martincorena *et al*, 2018). Furthermore, cancer genes are known to act in combination, and determining candidate sets of genes that are sufficient to cause cancer using genetic data alone would require astronomical sample sizes. Mechanistic studies are thus critical for conclusively determining that a particular gene is essential for cancer formation, and for identification of sets of genes that are sufficient for tumorigenesis.

In this work, we developed a novel cellular transformation assay that enables systematic testing of different combinations of oncogenic drivers in the context of cellular lineage. The strength of our approach is that we use human cells and known human oncogenes relevant to the specific cancer type in a molecularly defined assay that recapitulates the differentiation states between cell identities, creating a cellular state in which cells are susceptible to transformation (Fig. 6B). Recently, a similar approach using human pulmonary neuroendocrine cells derived from human embryonic stem cells (ESC) was used to generate xenograft tumors resembling small cell lung cancers by silencing retinoblastoma and *TP53* genes (Chen *et al*, 2019). Our assay differs from this important technology in that it avoids an intermediate cellular state that is tumorigenic (embryonic stem cells will form teratocarcinomas in mice). Furthermore, direct lineage conversion also allows more precise control of cell lineage than a differentiation protocol.

In the case of liver cancer, most evidence about the differentiated state required for transformation is based on mouse models. In mice, lineage-tracing has revealed that hepatocellular carcinoma initiates from mature hepatocytes, but sub-populations of tumor cells also show enrichment for stemness markers (Shin *et al*, 2016), supporting our finding that differentiation state can affect the susceptibility to cell transformation. *In vivo* in humans, a large subset of hepatocytes can re-enter the cell cycle after liver damage, potentially making them more susceptible to transformation (Fattovich *et al*, 2004). Consistently with this, in our assays, cultured non-proliferative hepatocytes were not transformed upon oncogene expression, and instead became senescent. When lineage-determining factors HNF1A, HNF4A, and FOXA3 were expressed together with the oncogenes, the senescence was blocked, but the cells still did not enter the cell cycle or transform. These results show that adult hepatocytes are resistant to oncogene expression and suggest that lineage-specific TFs might have a role in modulating this response. HNF1A, HNF4A, and FOXA3 are all expressed in human liver tumors at high–moderate levels as reported by TCGA (www.proteinatlas.org/), but their role in liver tumorigenesis is not yet clear. For example, tumor suppressive role has been suggested for HNF4A based on a rat model (Ning *et al*, 2010), but increased expression of a HNF4A transcript variant is associated with poor prognosis of HCC patients (Cai *et al*, 2017). Thus, the role of lineage-specific TFs and their expression level in liver tumorigenesis warrants further investigation.

Our approach allows a precise control of cell identity and differentiation state, facilitating analysis of interactions between driver genes, cell lineage and cell state (Fig. 6B). The results from our transformation model show that HFs can be directly converted to lineage-specific cancer by first inducing cell fate conversion towards hepatocyte identity with three lineage-specific TFs, HNF1A, HNF4A, and FOXA3, and then exposing the cells to liver cancer-specific oncogenes CTNNB1^T41A^, MYC, and TERT. We observed that all the oncogenes are necessary at the early stages of transformation. However, the mutant CTNNB1 was lost in the fully transformed tumorigenic iHeps whereas Wnt pathway modulator LGR5 was strongly up-regulated, demonstrating that our novel transformation assay can be used to study the dynamic features of the transformation process. However, comprehensive understanding of the mechanistic details still warrants further studies. Similarly, all three TFs are required for efficient lineage-conversion (Huang *et al*, 2014), but further experiments are necessary to identify the specific transcriptional mechanisms that enable the lineage-determining oncogenes to transform cells. This could, for example, be performed by testing combinations of different lineage-specific transcription factors with the oncogenes, and then identifying the specific target genes, and the promoter and enhancer elements required. Importantly, by using our novel cellular transformation assay, we were able to determine the minimum events necessary for making human liver cancer-like cells in culture. When transplanted to nude mice, they grow into tumors that are highly proliferative and malignant by morphology, and which would be classified as liver tumors based on expression of liver-specific markers used in differential diagnosis between sarcoma and hepatic tumors. By using lineage-specific TFs to generate the cell type of interest for transformation studies, our molecular approach can be generalized for identifying minimal determinants of any human cancer type, paving the way towards elucidating the exact molecular mechanisms by which specific combinations of mutations cause particular types of human cancer.

## Materials and methods

### Plasmids and lentiviral production

Full-length coding sequences for the TFs and oncogenes were obtained from GenScript and cloned into the lentiviral expression vector pLenti6/V5-DEST using the Gateway recombination system (Thermo Fisher Scientific). Expression construct for mCherry (#36084), lentiviral Cas9 expression construct LentiCas9-Blast (#52962), a cloning backbone lentiGuide-Puro (#52963), and the constructs for neuronal conversion (Tet-O-FUW-Ascl1, #27150; Tet-O-FUW-Brn2, #27151; Tet-O-FUW-Myt1l, #27152; Tet-O-FUW-NeuroD1, # 30129; pTetO-Ngn2-Puro, #52047; Tet-O-FUW-EGFP, # 30130; FUW-M2rtTA, #20342) were obtained from Addgene. The six pairs of single-stranded oligos corresponding to the guide sequences targeting the *TP53* gene in the GeCKO library were ordered from IDT, annealed, and ligated into lentiGuide-Puro backbone (Shalem *et al*, 2014). For virus production, the plasmids were co-transfected with the packaging plasmids psPAX2 and pMD2.G (Addgene #12260 and #12259, respectively) into 293FT cells (Thermo Fisher Scientific) with Lipofectamine 2000 (Thermo Fisher Scientific). Fresh culture medium was replenished on the following day, and the virus-containing medium was collected after 48 h. The lentiviral stocks were concentrated using Lenti-X concentrator (Clontech) and stored as single-use aliquots.

### Cell lines and generation of iHeps

Human foreskin fibroblasts (HFF, CCD-1112Sk) and human fetal lung (HFL) fibroblasts were obtained from ATCC (#CRL-2429 and #CCL-153, respectively) and cultured in fibroblast medium (DMEM supplemented with 10% FBS and antibiotics, Thermo Fisher Scientific). LentiCas9-Blast virus was transduced to early-passage fibroblasts (MOI = 1) with 8 µg/ml polybrene. Blasticidin selection (4 µg/ml) was started two days after transduction and continued for two weeks. Early passage blasticidin-resistant cells were used in the reprogramming experiments by transducing cells with constructs for TF expression in combinations reported earlier by Morris et al. (FOXA1, HNF4A, KLF5) (Morris *et al*, 2014), Du et al. (HNF4A, HNF1A, HNF6, ATF5, PROX1, CEBPA) (Du *et al*, 2014) and Huang et al. (FOXA3, HNF4A, HNF1A) (Huang *et al*, 2014) with MOI = 0.5 for each factor and 8 µg/ml polybrene (day 1). The medium was changed to fresh fibroblast medium containing β-mercaptoethanol on day 2 and to a defined hepatocyte growth medium (HCM, Lonza) on day On day 6, the cells were passaged on plates coated with type I collagen (Sigma) in several technical replicates, and thereafter, the HCM was replenished every two–three days.

Primary adult human hepatocytes (#HUCPI, batch HUM4122A, Lonza) were plated on type I collagen-coated 24-well plates in plating medium (MP100, Lonza) and maintained in hepatocyte growth medium (HCM, Lonza) as per vendor’s instructions. One day after plating, cells were transduced either with TFs that were used for iHep conversion (Huang *et al*, 2014) (FOXA3, HNF4A, HNF1A, MOI = 0.5) or with CMT oncogenes (CTNNB1^T41A^, MYC, TERT, MOI = 1) with 8 µg/ml polybrene in HCM medium. Fresh HCM medium was replenished on the following day and regularly thereafter. Seven days after iHep-TF transduction, these cells were transduced with CMT oncogenes as above. Beta-galactosidase staining for senescence analysis was performed three weeks after plating the cells.

Liver cancer cell lines HepG2 (#HB-8065, ATCC) and HuH7 (#JCRB0403, JCRB Cell Bank) were cultured in their recommended conditions and collected for gene expression analysis at ∼70% confluence. Culture medium for HepG2 comprises Eagle’s minimum essential medium supplemented with 10% FBS and antibiotics, and for HuH7 Dulbecco’s minimal essential medium with 10% FBS and antibiotics (Thermo Fisher Scientific).

### Generation of HCC-like cells

The iHeps generated using the three TFs (FOXA3, HNF4A, HNF1A) were passaged on type I collagen-coated plates on day 19 after iHep induction (p2) in HCM and transduced with different combinations of lentiviral constructs encoding the oncogenes (CTNNB1^T41A^, MYC, TERT, NFE2L2, PIK3CA^E545K^) on day 21 (MOI = 1 for each factor with 8 µg/ml polybrene). For CMT+sg*TP53* condition, the CMT oncogenes (CTNNB1^T41A^, MYC and TERT) were transduced along with a pool of six sgRNAs targeting the *TP53* gene. Fresh HCM was replenished on the day following the transduction, cells were maintained in HCM, and passaged when close to confluent. From fifth passaging onwards after oncogene induction, cells were maintained in HCM supplemented with 1% defined FBS (Thermo Fisher Scientific). For single-cell RNA-sequencing experiments, the iHeps were transduced with CMT oncogenes (MOI = 1 with 8 µg/ml polybrene) on day 8 with fresh HCM replenished on day 9, and the cells were harvested for single-cell RNA-sequencing at the indicated time points from replicate culture wells. In all experiments, viral construct for mCherry expression was co-transduced with the oncogenes. As controls, HFs were transduced with the same combination of oncogenes (CTNNB1^T41A^, MYC, TERT, MOI = 1 for each factor with 8 µg/ml polybrene). Fresh medium was changed on the day following the transduction, and the cells were passaged regularly. Cells were harvested for scRNA-seq analysis one and three weeks after transduction and used for beta-galactosidase staining three weeks after transduction.

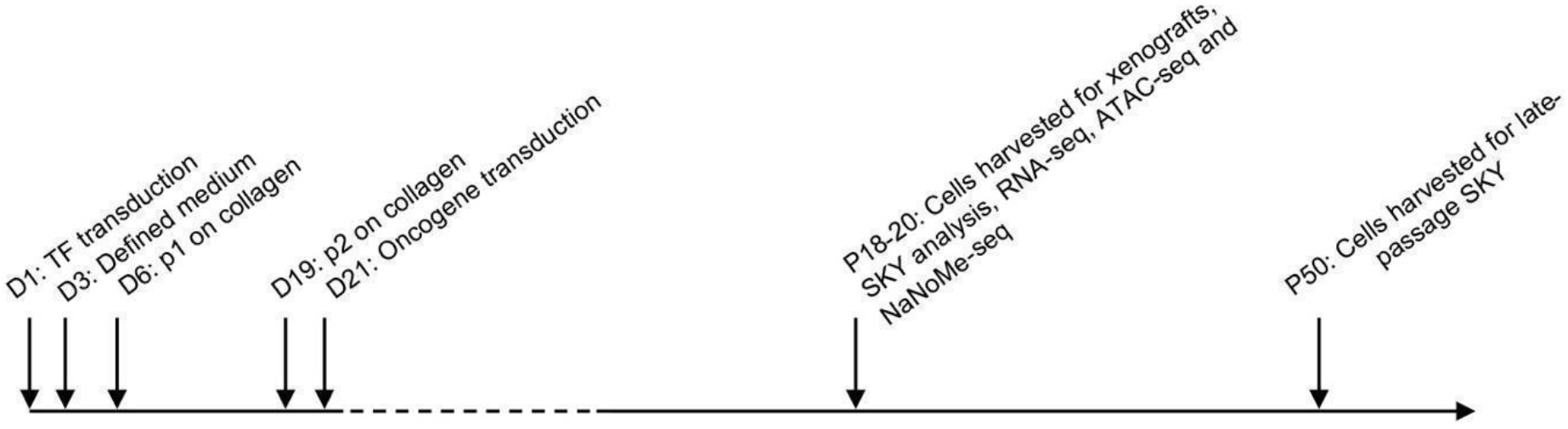

### Generation of induced neurons (iN)

HFs were plated on Matrigel-coated wells and transduced on the following day with tetracycline-inducible TF constructs for iN conversion (Ascl1, Brn2, Myt1l, NeuroD1, and Ngn2) with MOI = 0.3 for each factor and 8 µg/ml polybrene with co-transduction of lentiviral construct for EGFP; day 1). The medium was changed to fresh fibroblast medium containing β-mercaptoethanol on day 2. Doxycycline induction (2 ug/ml) was started on day 5, and the medium was replaced with defined N2B27 neuronal medium supplemented with small molecules (CHIR, SB431542, LDN, dcAMP, and Noggin) and doxycycline on day 6 and cells were maintained in the defined medium thereafter. At three weeks of iN conversion, cells were transduced with one of the two oncogene pools specific either for neuroblastoma (ALK^R1275Q^, MYCN, NRAS^Q61R^, PIK3CA^E545K^, BRAF^V600E^, PTPN11, PDGFRA, KIT, IDH1^R132H^) or for medulloblastoma (CTNNB1^T41A^, NRAS^Q61R^, PIK3CA^E545K^, SMO^W535L^, H3F3A^K28M^, IDH1^R132H^) with MOI = 0.5 for each factor and 8 µg/ml polybrene in neuronal medium. Fresh neuronal medium was replaced on the following day, the cells were maintained in neuronal medium and followed for 10-20 weeks.

### Xenografts

Oncogene-induced CMT and CMT+sg*TP53* cells at p20 (∼22-25 weeks after oncogene transduction) and control iHeps were harvested, 10^7^ cells were resuspended in HCM supplemented with 1% defined FBS and mixed with equal volume of Matrigel (growth factor reduced basement membrane matrix, Corning #356231) and injected subcutaneously into the flank of a 6-week old immunodeficient BALB/c nude male mice (Scanbur). Similarly, 10^7^ control fibroblasts were injected in equal volume of fibroblast medium and Matrigel. *In vivo* imaging of the tumors was performed for the mice under isoflurane anesthesia using the Lago system (Spectral Instruments Imaging). Photon counts from the mCherry were detected with fluorescence filters 570/630 nm and superimposed on a photographic image of the mice. Tumors were harvested 23-25 weeks after injection. All the experiments were performed according to the guidelines for animal experiments at the University of Helsinki and under license from appropriate Finnish Review Board for Animal Experiments.

### SKY analysis

Spectral karyotype analysis was performed at Roswell Park Cancer Institute Pathology Resource Network. Cells were treated for 3 hours with 0.06 µg/ml of colcemid, harvested and fixed with 3:1 methanol and acetic acid. Metaphase spreads from fixed cells were hybridized with SKY probe (Applied Spectral Imaging) for 36 hours at 37 degrees Celsius. Slides were prepared for imaging using CAD antibody kit (Applied Spectral Imaging) and counterstained with DAPI. Twenty metaphase spreads for each cell line were captured and analyzed using HiSKY software (Applied Spectral Imaging). In Fig. 3D, representative images are shown, and the recurrent chromosomal aberrations seen in majority (>90%) of the cells are reported.

### RNA isolation, qPCR and bulk RNA-sequencing

Total RNA was isolated from the control fibroblasts, liver cancer cell lines, iHeps harvested at day 5 and at weeks two, three, and four, CMT and CMT+sg*TP53* cells harvested at week two and week 22 (p20), and from tumor tissues stored in RNALater (Qiagen), using RNeasy Mini kit (Qiagen) with on-column DNase I treatment. For qRT-PCR analysis, cDNA synthesis from two biological replicates was performed using the Transcriptor High-fidelity cDNA synthesis kit (Roche) and real-time PCR using SYBR green (Roche) with primers specific for each transcript (Table S2). The Ct values for the target genes were normalized to those of GAPDH, and the mean values of sample replicates were shown for different conditions at the indicated time points. RNA-sequencing was performed from three biological replicate samples for each condition, using 400 ng of total RNA from each sample for poly(A) mRNA capture followed by stranded mRNA-seq library construction using KAPA stranded mRNA-seq kit for Illumina (Roche) as per manufacturer’s instruction. Final libraries with different sample indices were pooled in equimolar ratios based on quantification using KAPA library quantification kit for Illumina platforms (Roche) and size analysis on Fragment Analyzer (AATI) and sequenced on HiSeq 4000 (Illumina).

For preprocessing and analysis of the RNA-Seq reads the SePIA pipeline (Icay *et al*, 2016) based on the Anduril framework (Ovaska *et al*, 2010) was used. Quality metrics from the raw reads were estimated with FastQC (http://www.bioinformatics.babraham.ac.uk/projects/fastqc) and Trimmomatic (Bolger *et al*, 2014) clipped adaptors and low-quality bases. After trimming, reads shorter than 20 bp were discarded. Kallisto (v0.44.0) with Ensembl v85 (Zerbino *et al*, 2018) was used for quantification followed by tximport (Soneson *et al*, 2015) and DESeq2 (Love *et al*, 2014) (v1.18.1) for differential expression calculating log2(fold change) and standard error from triplicate samples. Human codon optimized lentiviral expression constructs were used for HNF4A, TERT and CTNNB1 (according to the recommended sequence by GenScript; Table S3). These codon-optimized sequences were included as additional transcripts in the reference genome for their identification from the RNA-sequencing data. Gene set enrichment analysis (Subramanian *et al*, 2005) was performed using GSEAPY (version 0.9.8) by ranking differentially expressed genes based on their −log10(p-value)*sign(fold-change) as metric. The gene signatures analysed for enrichment were collected from Molecular Signatures Database (MSigDB, version 6.2).

### Single-cell RNA-sequencing

For single cell RNA-sequencing (scRNA-seq), iHeps and HFs at different time points were harvested, washed with PBS containing 0.04% bovine serum albumin (BSA), resuspended in PBS containing 0.04% BSA at the cell density of 1000 cells / µl and passed through 35 µm cell strainer. Library preparation for Single Cell 3’RNA-seq run on Chromium platform (10x Genomics) for 4000 cells was performed according to manufacturer’s instructions and the libraries were paired-end sequenced (R1:27, i7-index:8, R2:98) on HiSeq 4000 (Illumina). Preprocessing of scRNA-seq data, including demultiplexing, alignment, filtering, barcode counting, and unique molecular identifier (UMI) counting was performed using CellRanger.

Quality control was applied separately for iHep and HFL-CMT samples. iHeps with fewer than 50,000 mapped reads or expressing fewer than 4000 genes or with greater than 6% UMI originating from mitochondrial genes were excluded, while for HFL-CMT samples, cells with fewer than 2500 genes or with greater than 10% UMI originating from mitochondrial genes were excluded. All genes that were not detected in at least 5 cells were discarded. From each sample, 500 cells were down-sampled for further analysis. The data was normalized and log-transformed using Seurat (Satija *et al*, 2015) (version 3.0.2). A cell cycle phase-specific score was generated for each cell, across five phases (G1/S, S, G2/M, M and M/G1) based on Macosko et al. (Macosko *et al*, 2015) using averaged normalized expression levels of the markers for each phase. The cell cycle phase scores together with nUMI and percentage of UMIs mapping to mitochondrial genes per cell were regressed out using a negative binomial model. The graph-based method from Seurat was used to cluster the cells. The first 30 PCs were used in construction of SNN graph, and 15 clusters were detected with a resolution of 0.8. Markers specific to each cluster were identified using the “negbinom” model. Pseudotime trajectories were constructed with URD (Farrell *et al*, 2018) (version 1.0.2). The RNA velocity analysis was performed using velocyto (La Manno *et al*, 2018) (version 0.17).

### Oil-Red-O- and PAS-staining and beta-galactosidase activity assay

Oil-Red-O and Periodic Acid-Schiff (PAS) staining were performed according to the manufacturer’s recommendation (Sigma). Briefly, for Oil-Red-O-staining, cells were fixed with paraformaldehyde (4%) for 30 mins, washed with PBS, incubated with 60% isopropanol for 5 mins and Oil-Red-O working solution for 10 mins, and washed twice with 70% ethanol. For PAS-staining, cells were fixed with alcoholic formalin (3.7%) for 1 min, incubated with PAS solution for 5 mins and Schiff’s reagent for 15 mins with several washes with water between each step, and counter-stained with hematoxylin. Beta-galactosidase assay was performed using Senescence detection kit (Abcam) according to the manufacturer’s protocol for fixation and staining (overnight).

### Immunohistochemistry

Tumor tissues were collected from mice injected with CMT+sgTP53 cells and fixed in 4% paraformaldehyde at 4°C overnight, dehydrated and embedded in paraffin. Five-μm sections were stained with hematoxylin and eosin with Tissue-Tek DRS automated system at Tissue Preparation and Histochemistry Unit (University of Helsinki, Finland) using standard protocols. For Ki-67 detection, sections were dewaxed with xylene, rehydrated, boiled in 10 mM citrate buffer, and treated with 3% hydrogen peroxide for 5 min to block the endogenous peroxidase activity. Sections were incubated with Anti-Ki-67 antibody (sc-101861, SantaCruz) at 4°C overnight followed by 40 min incubation with Brightvision Poly-HRP-Anti Mouse staining reagent (ImmunoLogic). The immune complexes were visualized using DAB Quanto chromogen and substrate (ThermoFisher) and counterstained with hematoxylin. The slides were dehydrated and mounted using Eukitt (Sigma). Tumor histology was analyzed by an experienced cancer pathologist.

### ATAC-seq

Fibroblasts and CMT+sg*TP53* cells (p20) were harvested and 50,000 cells for each condition were processed for ATAC-seq libraries using previously reported protocol (Corces *et al*, 2017) and sequenced PE 2×75 NextSeq 500 (Illumina). The quality metrics of the FASTQ files were checked using FASTQC and the adapters were removed using trim_galore. The reads were aligned to human genome (hg19) using BWA, and the duplicate reads and the mitochondrial reads were removed using PICARD. The filtered and aligned read files were used for peak calling using MACS2 and for visualizing the traces using the IGV genome browser.

### NaNoME-seq (NOME-seq using Nanopore sequencing)

To profile chromatin accessibility using GC methylase using NOME-seq protocol (Kelly *et al*, 2012) and to utilize the ability of Nanopore sequencing to detect CpG methylation without bisulfite conversion and PCR, we adapted the NOME-seq protocol for Nanopore sequencing on Promethion (NaNoME-seq). The nuclei isolation and treatment with GC methylase (M.CviPI) was performed as described earlier (Kelly *et al*, 2012) from the control fibroblasts and CHT+sgTP53 cells at p20. The DNA was isolated from GC methylase treated nuclei by phenol chloroform followed by ethanol precipitation. The sequencing library for Promethion was prepared using the 1D genomic DNA by ligation kit (SQK-LSK109) as per manufacturer’s recommendation and we loaded 50 fmol of final adapter-ligated high molecular weight genomic DNA to the flow cells for sequencing. After sequencing and base calling, the Nanopore reads were aligned to GRCh37 reference genome with minimap2 (Li, 2018). Nanopolish (Simpson *et al*, 2017) was modified to call methylation in GC context. In total, 11 Gbp of aligned read data from PCR amplified and GC methylated sequencing run was used to learn emission model for methylated GC sites. The learning process followed https://github.com/jts/methylation-analysis/blob/master/pipeline.make with adjustments for using human genome data and minimap2. For nuclear extract NaNoMe samples, methylation status was separately called for GC and CG sites. Similar independent method was recently described in a preprint by Lee et al (https://www.biorxiv.org/content/10.1101/504993v2). Reads with consecutive stretch of at least 80 GC sites with at least 75% methylated were filtered out due to expected cell free DNA contamination during library preparation as in Shipony et al. (https://www.biorxiv.org/content/10.1101/504662v1). The per site methylation levels in Fig. 5D are mean smoothed with triangular kernel 5 sites wide. Fibroblast and CMT+sg*TP53* NaNoMe analyses used 20.3Gbp and 24.8Gbp of aligned data, respectively.

Genomic reads from the NaNoMe-seq data were also used for detecting the number of insertions for the lentiviral expression constructs (**Table S1**). The lentiviral constructs were mapped to the nanopore reads and 1067 reads with lentiviral sequence were extracted, out of which 685 had some overlap with the inserted Cas9, CTNNB1, FOXA3, HNF1A, MYC or TERT constructs. Of these, 433 had at least 30bp alignment to both the lentiviral backbone and the insert. These reads were aligned to human genome, excluding alignments to native loci and locations with only one supporting read.

### Bisulfite-sequencing data

Publicly available bisulfite-sequencing data from the Roadmap Epigenomics Consortium was visualized at the marker gene promoters using Integrative Genomics Viewer. Wiggle files used were from human primary foreskin fibroblasts (GSM1127120_UCSF-UBC.Penis_Foreskin_Fibroblast_Primary_Cells.Bisulfite-Seq.skin03.wig.gz) and normal adult liver (GSM916049_BI.Adult_Liver.Bisulfite-Seq.3.wig.gz; GEO accession numbers GSM1127120 and GSM916049, respectively), representing methylation proportions [methylated calls / (methylated calls + unmethylated calls)] for all CpGs covered by at least 4 reads as documented in the GEO accession details.

## Acknowledgments

We thank Drs. Otto Kauko, Teemu Kivioja, and Minna Taipale for critical review of the manuscript, and Tomi Leung, Anu M. Luoto, Kaisu Jussila and Inga-Lill Åberg for technical assistance. We also thank HiLIFE research infrastructures including Biomedicum Virus Core, Single-cell Analytics FIMM, Biomedicum Imaging Unit and Laboratory Animal Center. We also thank the Roadmap Epigenomics Consortium for the publicly available data sets (http://nihroadmap.nih.gov/epigenomics/).

## Author contributions

JT conceived and supervised the study. BS designed and performed all the experiments with help from PP. BS performed the initial data processing of single cell and bulk RNA-seq data. KZ performed single cell RNA-seq analysis. BS performed the NaNoMe-seq and KP analyzed the data with inputs from SA and LA. AC and KZ performed bulk RNA-seq analysis with inputs from BS and PP. AR reviewed and analysed the immunohistochemistry slides, SH provided the bioinformatics support and inputs into the project. All authors contributed to the writing of the manuscript.

## Conflict of interests

Authors have no conflicts of interests.

## Corresponding author

Correspondence and requests for materials should be addressed to

## Funding

This work was supported by grants from Academy of Finland (Finnish Center of Excellence Program; 2012-2017, 250345 and 2018-2025, 312041, Post-doctoral fellowships; 274555, 288836 and Research Fellowships, 317807), Jane and Aatos Erkko Foundation, and the Finnish Cancer Foundation.

## Data and materials availability

All sequence data is available under ENA accession PRJEB31262.

## Notes

### Competing Interest Statement

The authors have declared no competing interest.

### Summary of Updates

Revised version with corrected text and figures

